# Incorporating neuronal fatigue in deep neural networks captures dynamics of adaptation in neurophysiology and perception

**DOI:** 10.1101/642777

**Authors:** Kasper Vinken, Xavier Boix, Gabriel Kreiman

## Abstract

Adaptation is a fundamental property of the visual system that molds how an object is processed and perceived in its temporal context. It is unknown whether adaptation requires a circuit level implementation or whether it emerges from neuronally intrinsic biophysical processes. Here we combined neurophysiological recordings, psychophysics, and deep convolutional neural network computational models to test the hypothesis that a neuronally intrinsic, biophysically plausible, fatigue mechanism is sufficient to account for the hallmark properties of adaptation. The proposed model captured neural signatures of adaptation including repetition suppression and novelty detection. At the behavioral level, the proposed model was consistent with perceptual aftereffects. Furthermore, adapting to prevailing but irrelevant inputs improves object recognition and the adaptation computations can be trained in a network trained to maximize recognition performance. These results show that an intrinsic fatigue mechanism can account for key neurophysiological and perceptual properties and enhance visual processing by incorporating temporal context.

## Introduction

Information processing in vision is not a constant function of visual input, but rather the underlying computations are dynamically changed by the behavioral and temporal context. These dynamic changes can dramatically alter visual experience, such as the illusory perception of upwards motion after watching flowing water in the so-called waterfall illusion^1^. To understand vision under natural, dynamic conditions, we must consider the processes that contribute to the integration of temporal context.

A particularly notable aspect of how temporal context is incorporated in visual processing is the effect of adaptation on neural activity and perception. Adaptation generally refers to the dependence on recent stimulation and is considered to be an evolutionary conserved and fundamental property of visual and other sensory systems^2^. Perceptually, exposure to a stimulus often leads to a temporarily reduced sensitivity for its features^3^. The lingering effects after removal of the stimulus are called aftereffects, which have been described for a wide range of low to high-level visual stimulus properties (for an overview, see^4^). Likewise, exposure to an adapter stimulus often reduces the neural response to a subsequent test stimulus that is similar to the adapter. The dependence on the similarity between adapter and test stimulus (i.e. stimulus specificity), is a hallmark property of adaptation (for an overview, see^5^). When a stimulus is repeated, the resulting response reduction is called repetition suppression. Because of their conceptual similarity, repetition suppression and aftereffects are usually considered to be manifestations of the same underlying mechanisms; yet, those mechanisms remain poorly understood.

The dynamics of adaptation could be entirely implemented by circuit-level interactions in the neural network, but they could also arise from intrinsic biophysical mechanisms in each individual neuron^2^. For example, the responsiveness of a cortical neuron can be controlled by mechanisms that increase the membrane conductance. Indeed, contrast adaptation in cat visual cortex leads to a strong afterhyperpolarization of the membrane potential^6,7^. This afterhyperpolarization is caused by sodium-activated potassium currents that are triggered by the influx of sodium ions following high frequency firing^8,9^. Thus, in this scenario, intrinsic properties of individual neurons control their responsiveness based on the strength of their previous activation, a phenomenon referred to as firing-rate adaptation or fatigue. Neuronal fatigue on its own cannot explain complex effects of adaptation such as stimulus selectivity, because fatigue should equally affect responses for all stimuli. However, several studies have suggested that fatigue effects cascade through the visual system^10,11^ and can influence downstream processing in unexpected ways^2,12^.

In this study, we evaluate whether an intrinsic neuronal fatigue mechanism can account for complex key neurophysiological and perceptual adaptation phenomena in the visual system. To evaluate this hypothesis, we build artificial neural network models and compare their outputs with both neurophysiological recordings and behavioral measurements. We focus on a family of feedforward artificial neural networks that has been shown to provide a reasonable first-order approximation to describe ventral visual stream responses to brief stimulus presentations^13–18^ while capturing aspects of object recognition and perceived shape similarity^14,17,19^. These networks are based on hierarchical cascades of simplified linear summation units plus a nonlinear activation function, and do not incorporate the temporal dynamics of intrinsic neuronal mechanisms. Implementing these dynamic mechanisms can provide a comprehensive framework for connecting the different levels at which adaptation phenomena have traditionally been described, which is an important step towards understanding the role of adaptation in information processing.

We started by implementing an exponentially decaying fatigue mechanism in each unit of a deep neural network pre-trained on object recognition^20^. This model could account for key temporal dynamics of adaptation in neurophysiology and perception. Next, using a psychophysics experiment, we show that adapting to prevailing conditions can lead to improved object recognition in both human participants as well as the proposed computational model. Furthermore, adaptation could be optimized in a network trained to maximize recognition performance in the same psychophysics task. Finally, we show that a circuit solution learned by a recurrent neural network is less robust than intrinsic neuronal fatigue.

## Results

The goal of this study was to examine whether intrinsic neuronal fatigue can account for adaptation phenomena in the visual system. We introduced an adaptive mechanism into bottom-up computational models of vision by implementing units that show neuronal fatigue (**Fig. 1**, Methods). We show that the incorporation of neuronal fatigue is able to capture fundamental dynamic properties of adaptation both at the neurophysiological level as well as at the behavioral/perceptual level.

**Fig. 1.**
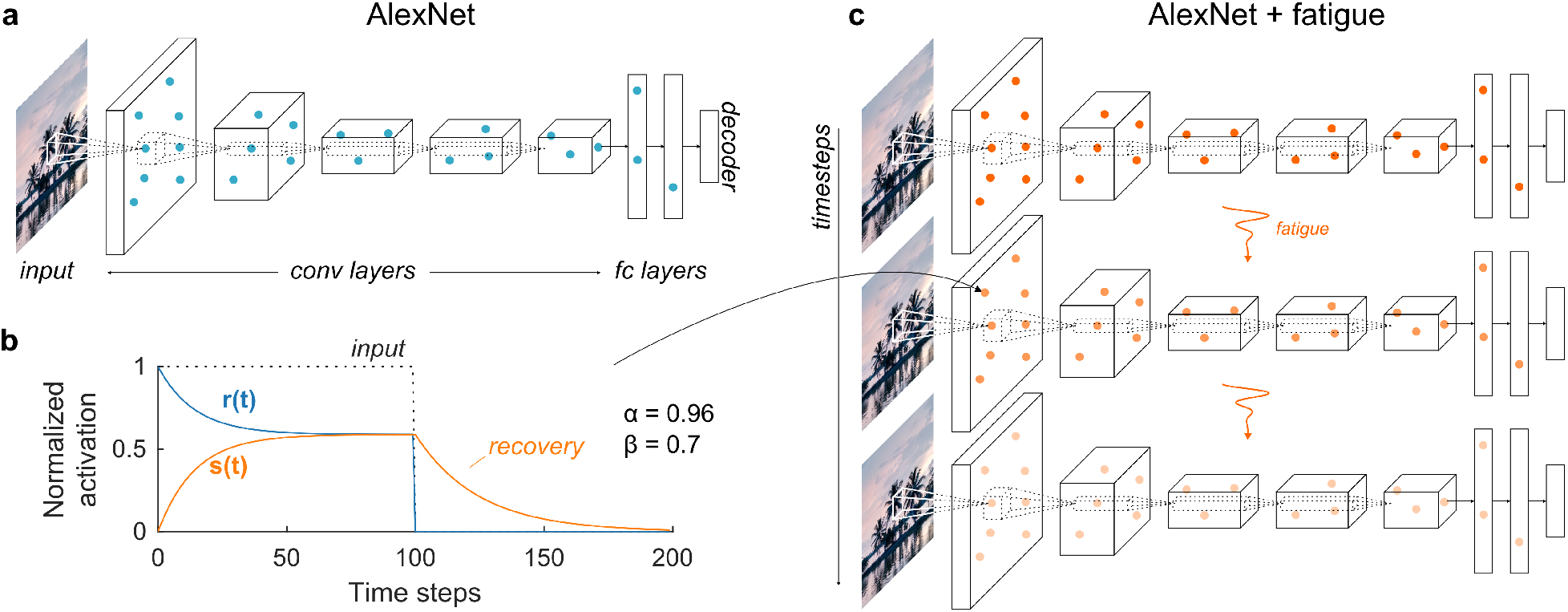
Neural network architecture and incorporation of intrinsic neuronal fatigue mechanism. **a**, Architecture of a static deep convolutional neural network, in this case AlexNet^20^. AlexNet contains five convolutional layers (conv1-5) and three fully connected layers (fc6, fc7, and the decoder fc8). The unit activations in each layer, and therefore the output of the network, are a fixed function of the input image. **b**, Fatigue implemented by Equations 1 and 2 results in suppression *s*(*t*) (orange) over time for constant input (time steps 0-100), leading to a reduction in the response *r*(*t*) (blue). The amount of suppression recovers in the absence of input (time steps > 100). In this case, the parameters in 1 and 2 take the values *α* = 0.96 and *β* = 0.7. **c**, An expansion over time of the network in **a**, where the activation of each unit is a function of its inputs *and* its activation at the previous time step (Equations 1 and 2).

### A neural network incorporating neuronal fatigue captures the attenuation in neurophysiological responses during repetition suppression

The most prominent characteristic of neural adaptation is repetition suppression, a reduction in the neuronal responses when a stimulus is repeated. For example, in an experiment with sequential presentation of two stimuli, the response to the second stimulus is typically lower and this reduction is strongest when the second stimulus is identical to the first one. We illustrate this phenomenon using an experiment where face stimuli lasting 250 milliseconds were presented to a macaque monkey in separate repetition trials (the same face repeated) and alternation trials (two different faces shown), with an interval of 500 milliseconds in between stimuli (**Fig. 2a**,^21^). The monkey was performing an unrelated task: detecting catch trials consisting of an inverted face (the neural responses to the inverted face trials are not shown here). Neurons recorded in the middle lateral face patch of inferior temporal (IT) cortex showed a decrease in the response during stimulus presentation and after stimulus offset. In addition, the neurons showed a lower response to a face stimulus when it was a repetition trial (blue) compared to an alternation trial (orange; **Fig. 2b**). It is not straightforward to see how neuronal fatigue could account for stimulus-specific repetition suppression, because an activation-based adaptation mechanism by itself is not stimulus selective. Selectivity could emerge in a network where different stimuli excite and cause fatigue in not entirely overlapping input populations ^23^. To evaluate this possibility, we implemented a neuronal fatigue mechanism (**Fig. 1b**) into the responses of units of AlexNet^20^ (**Fig. 1a**), an image-computable deep convolutional neural network model (Methods). Due to the fatigue mechanism, the model units display temporally evolving responses (**Fig. 1c**) and can be directly compared to the neurophysiological dynamics. Like the biological neurons, the model units also showed a decrease in the response during the course of stimulus presentation. Also consistent with the biological responses, we found stimulus-specific repetition suppression when we presented the same stimuli shown to the monkeys (**Fig. 2c**, average of all *N* = 6,699 units with a positive net response to faces in layer conv5). We focus here on the outputs of the conv5 layer (after the rectification and linearization operation, ReLU), which is presumed to show a stronger correlation to macaque IT responses than other layers^24^. In addition, for this experiment, units with a positive net response from layer conv5 showed the largest average stimulus-specific suppression. The model units demonstrate the key features of adaptation at two time scales: (i) during presentation of any stimulus, including the first stimulus, there is a decrease in the response with time; (ii) the overall response to the second stimulus is smaller than the overall response to the first stimulus; (iii) the response to the second stimulus is attenuated more when it is a repetition.

**Fig. 2.**
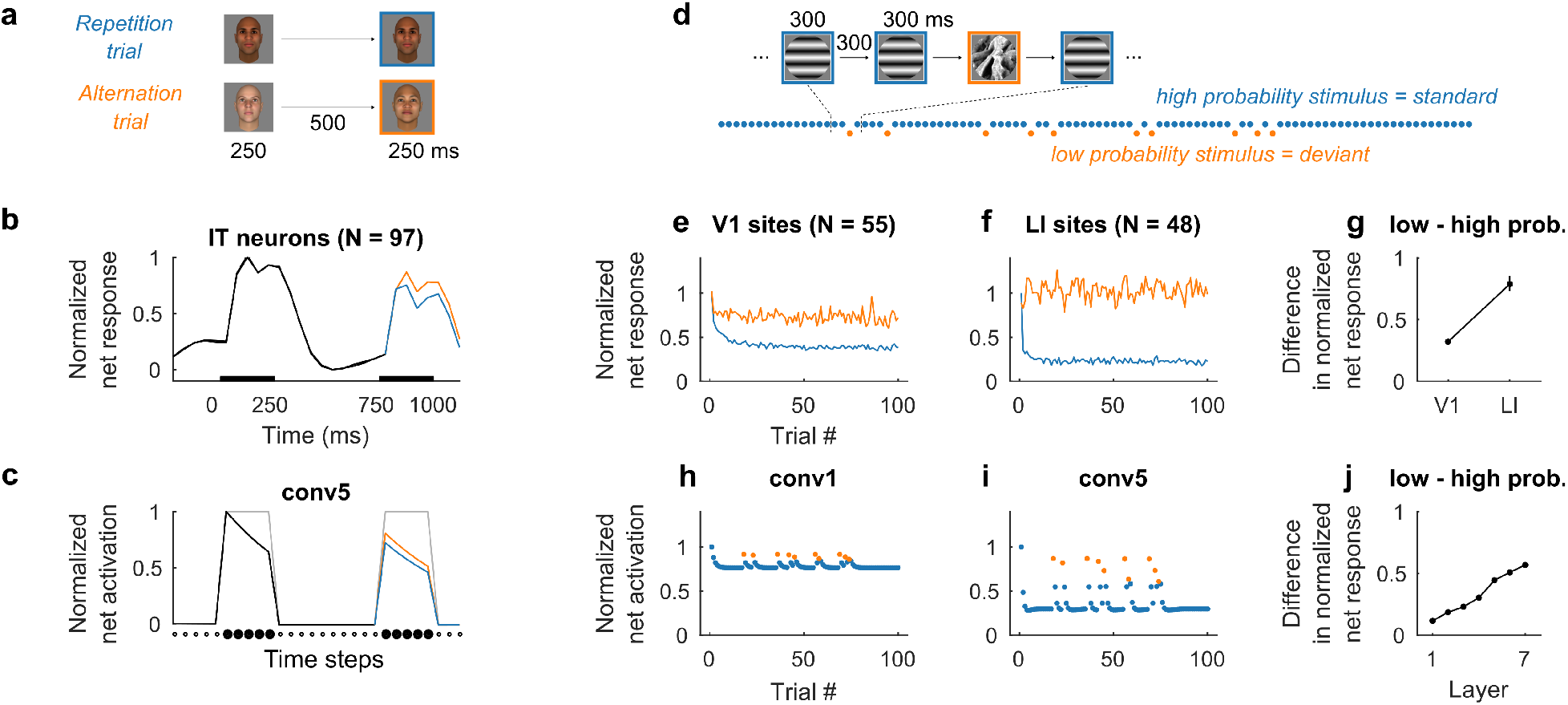
Neuronal fatigue in a neural network captures temporal dynamics of adaptation at the neurophysiological level. **a**, Example repetition and alternation trials. **b**, Responses in macaque inferior temporal (IT) cortex (*N* = 97 neurons with a positive net response to faces) are suppressed more for a repeated stimulus (blue) than for a new stimulus (orange, data from^21^). Black bars indicate stimulus presentation. **c**, The same experiment as in **a-b** leads to stimulus-specific suppression similar to the experimental results in an artificial neural network with neuronal fatigue (black, blue and orange lines; average activity after ReLU of *N* = 6,699 conv5 units with a positive net response to faces). Suppression is not present without neuronal fatigue (grey). The x-axis units are time steps, mapping to bins of 50 ms in **b. d**, Example oddball sequence with illustrative stimuli: a high probability grating (blue) and a low probability texture (orange) used in the actual and simulated experiments. Stimulus type and probability were counterbalanced for each neural recording. **e,f**, Neural responses from multi-unit sites recorded during oddball sequences in rat primary (V1, *N* = 55, e) and latero-intermediate (LI, *N* = 48, **f**) visual cortex^22^. Accumulation of adaptation across multiple repeats leads to increased suppression for high probability stimuli (blue) compared to low probability stimuli (orange). Time courses were normalized by the response at the first trial. **g**, Difference in response for the low and high probability stimulus increases from V1 to LI (error bars are 95% bootstrap confidence intervals calculated assuming no inter-animal difference). **h-j**, A simulation of an oddball sequence shown in **d** in the computational model leads to similar accumulation of suppression across stimulus presentations and stages of processing as in the experimental results.

In addition to the two temporal scales illustrated in **Fig. 2a-c**, adaptation not only affects responses from one stimulus to the next, but also operates at longer time scales. For example, repetition suppression typically accumulates across multiple stimulus presentations and can survive intervening stimuli^25^. To illustrate this longer time scale over multiple trials, we present data from rat visual cortex^22^ during an *oddball paradigm* where two stimuli are presented in a sequence with different probabilities (**Fig. 2d**): the *standard* stimulus is shown with high probability (blue) and the *deviant* stimulus is shown with a low probability (orange). The sequence consisted of 100 stimulus presentations, each one shown for 300 ms and separated by 300 ms, with 90 standard stimuli and 10 deviant stimuli shown in random order. The standard stimulus is far more likely to be repeated in the sequence, allowing adaptation to build up and therefore causing a decrease in the response for later trials in the sequence (**Fig. 2e,f**, blue). This observation is evident both in primary visual cortex (V1) and in the extrastriate latero-intermediate visual cortex (LI). In contrast, the low probability stimulus does not show such a response reduction (**Fig. 2e,f**, orange). The proposed computational model, the same model used in **Fig. 2a-c** without any tuning or changes, was able to qualitatively capture this response difference between standard and deviant stimuli (**Fig. 2h,i**). The model also showed a partial release from adaptation after each presentation of a deviant stimulus (i.e., a slight increase in the response to the next standard stimulus following presentation of a deviant stimulus), similar to the observations in rat visual cortex (see Supplemental Information of ^22^). The LI responses in **Fig. 2f** also showed an *increased* response to the low probability deviant (orange normalized net response above 1). This effect was not captured by the model and is likely to require additional mechanisms (see Discussion).

An additional observation from these data is the increase in deviant-standard stimulus response difference moving from V1 to LI **Fig. 2g**. The increase in adaptation along the visual hierarchy is consistent with the idea of adaptation cascading through the visual system, with additional contributions at multiple stages. This increase was also reflected in the proposed computational model where adaptation effects increased from one layer to the next (**Fig. 2j**). Together, these results demonstrate that neuronal fatigue propagating through the visual hierarchy leads to encoding of stimulus occurrence probabilities at multiple timescales, with increasing sensitivity ascending through the ventral visual stream.

### A neural network incorporating neuronal fatigue reveals perceptual manifestations of adaptation

A comprehensive model of visual adaptation should not only capture the neural properties of repetition suppression, but should also be able to explain perceptual aftereffects of adaptation. We start with the well-known tilt aftereffect^26,27^, where an observer perceives a vertically oriented grating to be slightly tilted in the direction *opposite* to the tilt direction of an adapter. In other words, the decision boundary for perceptual orientation discrimination shifts towards the adapter^27^, and a grating tilted slightly in the direction of the adapter will appear vertical (see **Fig. 3a**). No perceptual shift should occur when the adapter has the same orientation as the original boundary stimulus.

**Fig. 3.**
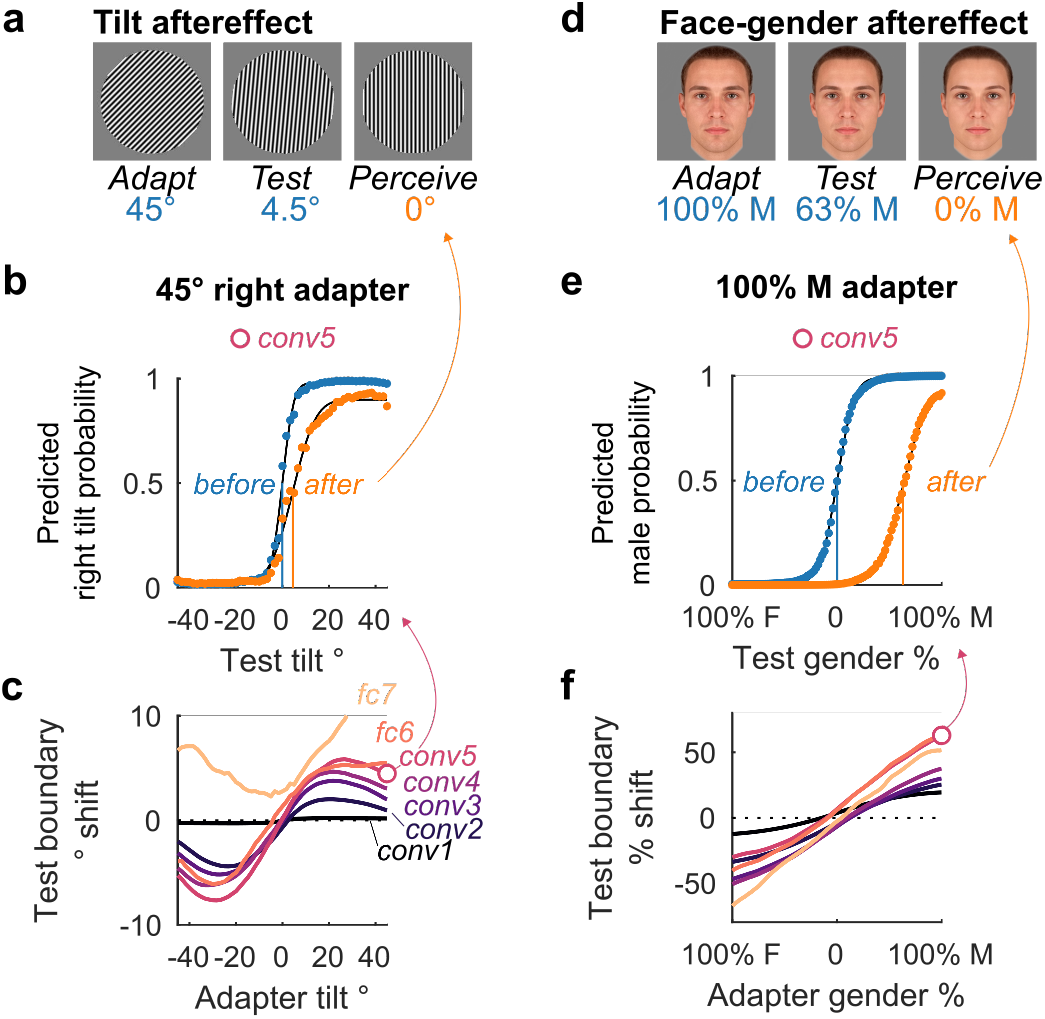
A neural network incorporating neuronal fatigue demonstrates perceptual adaptation effects. **a,d**, Illustration of the tilt (**a**) and face-gender (**d**) aftereffect with the stimuli used in our simulated experiments. After exposure to an adapter (left), a test stimulus is presented (middle). The test stimulus is perceived differently as a result of a shift in the decision boundary toward the adapter. In **a**, observers perceive the slightly right tilted grating as vertically oriented. In **d**, observers perceive the male face as gender-neutral. The images for the illustration of perceived aftereffects (**a,d**) were picked based on the estimated conv5 boundary shift shown below. **b,e**, Decision boundaries before (blue) versus after (orange) exposure to the adapter based on the conv5 layer of the model with neuronal fatigue. Markers show class probabilities predicted by the logistic regression classifier for each test stimulus, full lines indicate the corresponding psychometric functions, and vertical lines the classification boundaries. In **b**, adaptation to a 45° grating leads to a shift in the decision boundary to positive orientations, hence perceiving test stimuli to the left of the new boundary with more negative orientations. In **e**, adaptation to a 100% male face leads to a shift in the decision boundary towards male faces, hence perceiving test stimuli to the left of the new boundary as more female-like. **c,f**, Decision boundary shifts for the test stimulus as a function of the adapter tilt/face-gender per layer. Round markers indicate the conv5 boundary shifts plotted in **b,e**.

To evaluate whether perceptual aftereffects can be described by the proposed neural network model with fatigue, we created a set of gratings that ranged from left to right oriented (−45° to 45° in 100 steps), and measured the category boundary for each layer of the model before and after adaptation. Specifically, these boundaries were estimated using a binary classifier (logistic regression, trained on the full stimulus set before adaptation) and fitting a psychometric function^28^ on the class probability estimates given by that classifier. For a fair comparison, the categorical boundaries before and after adaptation were always calculated using the same classifier in the same space, namely the principal component space obtained from the unadapted outputs to the full stimulus set. Of note, the model used here is the same one used in **Fig. 2**. In **Fig. 3b** we show the psychometric curves fit on the conv5 layer class probability estimates before (blue) and after (orange) adaptation to a 45° right tilted grating. The decision boundary, that is the test stimulus tilt for which the classifier’s predicted right tilt probability was 0.5, shifted 4.5° towards the tilt of the adapter. **Fig. 3c** shows that for all layers except fc7, adaptation to a tilted grating resulted in a boundary shift towards the adapter. Given that the original boundary stimulus is vertical, adaptation to a vertically oriented grating had no effect on the decision boundary. The effect of adaptation propagated and accumulated over the layers, and the shift in the test boundary was largest for conv5 and fc6. For conv1 there was also a shift in the predicted direction, which is too small to notice at this scale in **Fig. 3c**.

Aftereffects have also been described for more complex stimulus properties, such as the gender of faces^29^. Like the tilt aftereffect, adapting to a male or female face results in a face-gender boundary shift towards the adapter, whereas adapting to a gender-neutral face leads to no shift. In other words, exposure to a male face will make a face appear more female (**Fig. 3d**). To evaluate whether the model also shows the face gender aftereffect, we created a set of face stimuli that morphed from average female to average male face in 100 steps (using Webmorph^30^). As predicted, exposing the model to an adapter face shifted the face gender boundary towards the adapter gender (**Fig. 3e,f**). For example, before adaptation, the predicted male probabilities for the model conv5 layer showed a typical sigmoidal curve centered around the gender-neutral face stimulus (**Fig. 3e**, blue). After adapting to a 100% male face, the decision boundary (i.e. predicted male probability of 0.5) shifted 63 percentage values towards the gender of the adapter (**Fig. 3e**, orange). As predicted, adaptation to a gender-neutral face had no effect (**Fig. 3f**, 0% on the x-axis). The aftereffect did not suddenly emerge in later layers, but slowly built up in an approximately monotonic fashion with increasing layers (**Fig. 3f**, from black to blue to red colors), consistent with the idea of adaptation cascading and accumulating across the stages of processing. All stages of processing can contribute to the aftereffects, based on the population of units that responds to both the adapter and test images.

### Adapting to prevailing but irrelevant input enhances object recognition

One of the proposed computational roles of neural adaptation is to decrease salience of recently seen stimuli or features^12,23^ and increase sensitivity to small changes in the sensory environment^5,32^. To test this hypothesis in an object recognition task, we developed a psychophysics experiment where participants were required to classify hand drawn doodles (**Fig. 4a**) hidden in a noise pattern. We asked whether adaptation to a noise pattern would increase the ability to recognize the target object embedded in the same noise pattern. We compared the behavioral results against two control conditions where we would expect to see no effect of adaptation: one where no adapter was used (i.e. an empty frame, **Fig. 4b**, left) and one where a different noise pattern was used as adapter (**Fig. 4b**, right, see Methods).

**Fig. 4.**
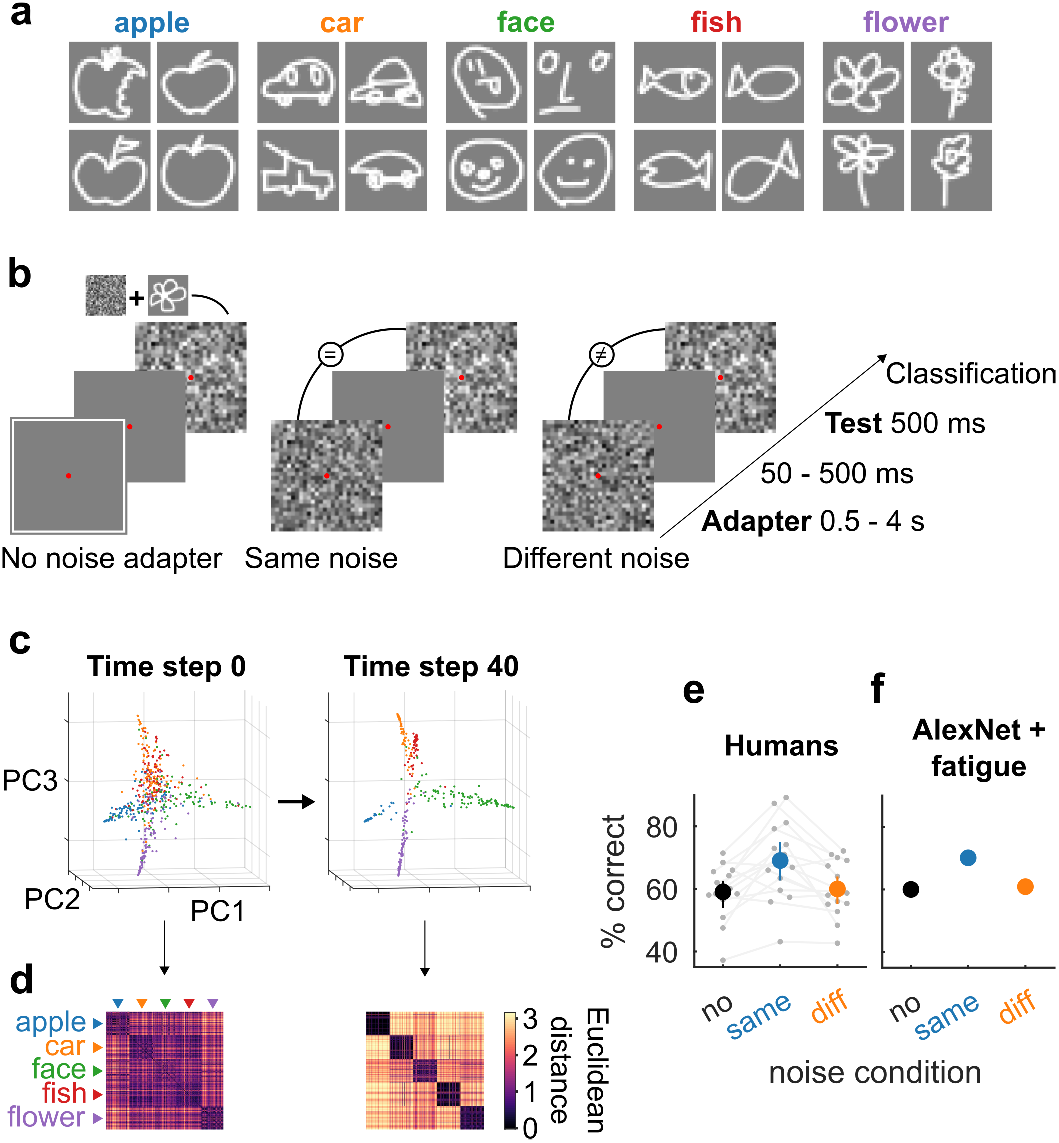
Neuronal fatigue increases sensitivity to changes by adapting to previous input. **a**, Representative examples for each of the five doodle categories from the total set of 540 selected images^31^. **b**, Schematic illustration of the conditions used in the doodle experiment. In each trial participants or the model had to classify a hand drawn doodle hidden in noise (test), after adapting to the same (middle), a different (right), or no (left) noise pattern. **c,d**, Neuronal fatigue in the proposed computational model enhanced the representation of the signal (doodle) by adapting to the prevailing sensory conditions (noise pattern). **c**, Dynamic evolution of the representation of the images embedded in noise while adapting to the same noise pattern for 40 time steps. The 3 axes correspond to the first 3 principal components of the fc8 layer representation of all the test images. Each dot represents a separate doodle+noise image, the color corresponds to the category (as shown by the text in **d**). Adaptation to the same noise pattern moves the doodle representations in fc8 principal component space to separable clusters. **d**, Dissimilarity matrix for all pairs of images. Entry (i,j) shows the Euclidean distance between image i and image j based on the fc8 features at time step 0 (left) or time step 40 (right). The distance is represented by the color of each point in the matrix (see scale on right). Images are sorted based on their categories. Adaptation leads to an increase in between category distances and a decrease in within category distances as shown by the pairwise distance matrices. **e,f**, Both human participants (**e**) and the proposed computational model (**f**) showed an increase in categorization performance after adapting to the same noise pattern. Gray circles and lines denote individual participants (*N* = 15). The colored circles show average categorization performance, error bars indicate 95% bootstrap confidence intervals. Chance = 20%. Note that for **f** we decreased the suppression scaling constant to *β* = 0.1 to get comparable adapter effects (for all other figures it was set to *β* = 0.7).

We first evaluated the effect of adaptation on the representations of test images in the proposed computational model for the same-noise condition. **Fig. 4c** shows the temporal evolution of the representation of each noisy doodle in a space determined by the first 3 principal components of the fc8 outputs: adaptation to the noise pattern moves the colored dots representing doodle images into distinctly separable clusters. We further quantified this separation in feature space by computing the dissimilarity matrix for all possible pairs of images (**Fig. 4d**). Adaptation led to increased differentiation of the between-category comparisons (off diagonal squares) and increased similarity between images within each category (diagonal squares) from the initial conditions (left) to the final time step (right).

Next, we evaluated whether adaptation could increase the ability of human participants to recognize doodles embedded in noise. Recognizing these doodles in this task is not trivial: whereas subjects can readily recognize the doodles in isolation, when they are embedded in noise and in the absence of any adapter, categorization performance was 59% (*SD* = 8.7%) where chance is 20%. Adapting to the same noise pattern increased categorization performance by 10% (**Fig. 4e**, *p* = 0.0026, Wilcoxon signed rank test, *N* =15 subjects). This increase in categorization performance was contingent upon the noise pattern presented during the test stimulus being the same as the noise pattern in the adapter. Performance in the same-noise condition was 9% higher than in the different-noise condition (*p* = 0.0012, Wilcoxon signed rank test, *N* =15 subjects). We next evaluated the model’s performance in this same task, after fine-tuning the fully connected layers to classify doodles (see Methods). The model demonstrated the same effects as the human participants, showing increased performance for the same-noise condition compared to the no adapter condition or different-noise condition (**Fig. 4f**). Thus, adapting to a prevailing noise pattern improved the ability to recognize test images and this effect can be accounted for by neuronal fatigue in a feed-forward neural network.

### Adaptation parameters can be trained by maximizing recognition performance

The parameters *α* and *β* for the adaptation state considered thus far were chosen to impose neuronal fatigue. Rather than choosing those parameters, here we asked whether it is feasible to fit them in the doodle experiment described in the previous section using recognition performance as the goal. In order to test whether neuronal fatigue can emerge from training, we made a smaller network with an AlexNet-like architecture (but without response normalization) consisting of three convolutional and two fully connected layers (see **Fig. 5a**, without the recurrent connections shown in blue, which are discussed in the next section). Each unit (excluding the decoder layer) had an exponentially decaying adaptation state as defined by Equations 1 and 2. For simplicity, the trials of the doodle experiment were presented in only three time steps: the adapter at step one, a blank interval at step two, and the test image at step three (**Fig. 5a**). In addition to training the feedforward weights on the task, we also trained separate *α* and *β* parameters per layer. The value of *α* determines how fast the hidden state updates, ranging from no update (*α* = 1) to completely renewing at each time step (*α* = 0). The value of *β* determines whether the hidden state is used for activation-based fatigue (*β* > 0), enhancement (*β* < 0) or nothing at all (*β* = 0). After training using 30 random initializations of the network on same-noise trials, the resulting parameters revealed activation-based fatigue which was particularly strong for convolutional layers 1 and 2 as indicated by the *positive* high *β* and relatively low *α* values (**Fig. 5b**). The average categorization performance on the test set was 97.9% (blue), compared to 74.8% when no adaptation state was included (black; **Fig. 5c**). This categorization performance was much higher than the numbers reported in **Fig. 4f**, where the parameters *α* and *β* were fixed rather then optimized for this specific task. When the network was trained and tested on different-noise trials, categorization performance on the test set was similar to the one obtained without adaptation state (orange; **Fig. 5c**). Thus, when a network with adaptation state is trained on an object recognition task with a temporally constant but irrelevant input pattern, the optimized adaptation parameters show activity-based suppression for its units.

**Fig. 5.**
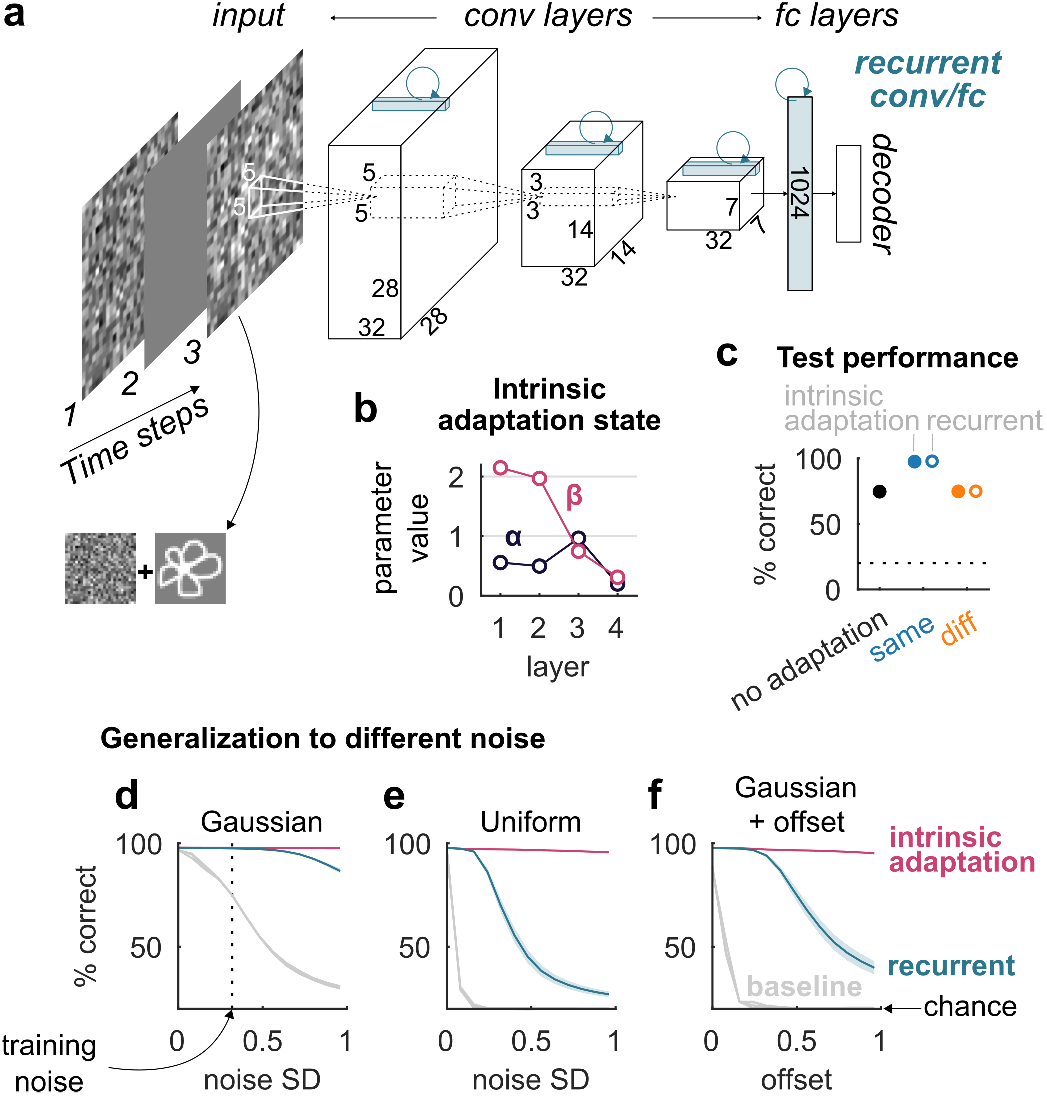
The parameters for adaptation can be fit in a network trained to maximize recognition performance and this solution is robust to different input signals. **a**, A convolutional neural network with an AlexNet-like feed-forward architecture. For the adaptation version, an exponentially decaying hidden state was added to each unit according to Equations 1 and 2 (except for the decoder). For the recurrent version, fully connected recurrent weights were added for the fully connected layer and convolutional recurrent kernels of size 1 × 1 × 32 and stride 1 for the three convolutional layers (see drawings in blue). **b**, Average fitted parameters *α* and *β* for each layer after training 30 random initializations of the network with adaptation state on same noise trials (standard error of the mean bars are smaller than the markers). **c**, Test categorization performance on trials with the same Gaussian noise distribution as during training. Full markers: average categorization performance after training 30 random initializations on the same noise trials without adaptation state (black), after training with adaptation state on same noise trials (blue) or on different noise trials (orange). Empty markers: same as full markers but for the recurrent neural network. Standard error of the mean bars are smaller than the markers. Chance = 20%, indicated by horizontal dotted line. **d-f**, Average generalization performance of the networks with an intrinsic adaptation state (magenta), recurrent weights (blue), or neither (grey, baseline) for same noise trials under noise conditions that differed from training. Performance is plotted as a function of increasing standard deviations (x-axis) of Gaussian noise (**d**, the vertical line indicates the SD = 0.32 used during training), of uniform noise (**e**), or as a function of increasing offset values added to Gaussian noise (SD = 0.32, same as training). Error bounds indicate standard error of the mean.

### Intrinsic fatigue generalizes better to different input pattern distributions than a circuit solution

A common way to model temporal dynamics in the visual system is by adding recurrent weights to a feedforward network^33,34^, and recurrent neural networks can demonstrate phenomena similar to adaptation^35^. Recurrent neural networks are the standard architectures used to process input sequences and should be able to perform well in the noisy doodle categorization task. To compare the intrinsic fatigue mechanism with a circuit solution, we considered a network without an exponentially decaying adaptation state and added lateral recurrent connections illustrated in blue in **Fig. 5a**. After training on same-noise and different-noise trials, the recurrent architecture achieved the same categorization performance on the test set as the architecture with an adaptation state (**Fig. 5c**). Thus, as expected, the recurrent network performed on par with the network with trained fatigue when the same Gaussian noise distribution was used during training and testing. To further investigate the two possible solutions, we considered situations where the distribution of noise patterns used during training and testing was different. We reasoned that a trained intrinsic fatigue mechanism should generalize well across different input features or statistics, whereas the circuit-based solution learned by a recurrent neural network might be less robust. In contrast with the intrinsic mechanism, the recurrent neural network requires a large number of parameters for its recurrent loops to store and subtract previous input, and might therefore over-fit to the particular input statistics of the adapter during training. Indeed, the recurrent network failed to generalize to higher standard deviations of Gaussian noise (**Fig. 5d**), and failed dramatically when tested with uniformly distributed noise (**Fig. 5e**), or Gaussian noise with an offset (**Fig. 5f**). In stark contrast, the intrinsic fatigue mechanism generalized well across all of these different input noise changes (**Fig. 5d-f**, magenta). In sum, while a recurrent network circuit implementation can learn to solve the same task, the solution is not robust to deviations from the particular statistics of the prevailing input during training.

## Discussion

We examined whether the paradigmatic neurophysiological and perceptual signatures of adaptation effects can be explained by a biologically inspired intrinsic fatigue mechanism^7^ in a feed-forward deep network. The proposed computational model bridges the fundamental levels at which adaptation phenomena have been described: from intrinsic cellular mechanisms, to responses of neurons within a network, to perception. We showed that the proposed model (**Fig. 1**) can explain classical perceptual aftereffects of adaptation, such as the tilt and face-gender aftereffects^26,29^ (**Fig. 3**). In addition, the units in this network exhibited stimulus-specific repetition suppression^5,23^, which recovers over time but also builds up across repeats despite intervening stimuli^36^, and builds up across stages of processing^22,37,38^ (**Fig. 2**). Such adaptation effects may enhance novelty detection. In addition, adaptation can directly impact recognition performance (**Fig. 4**). As predicted by the proposed model, adapting to a noise pattern increased object recognition performance when the target object was hidden in a temporally constant noise pattern. A recurrent neural network could be trained to solve the same task, but it converged on a circuit solution that was less robust to different noise conditions than the proposed model based on intrinsic neuronal fatigue mechanisms (**Fig. 5**). Together, these results show that a neuronally intrinsic fatigue mechanism can robustly account for adaptation effects at the neurophysiological and perceptual levels.

Adaptation can serve an important metabolic role. Neurons use large amounts of energy to generate action potentials, which constrains neural representations^39,40^. By reducing neural responsiveness for redundant information, adaptation has the advantage of increasing metabolic efficiency of the neural code. Such metabolic advantages could also serve to increase coding efficiency by normalizing responses for the current sensory conditions^5^. Neurons have a limited dynamic range with respect to the feature they encode and a limited number of response levels. Adaptation can maximize the information carried by a neuron by re-centering tuning around the prevailing conditions and thus preventing response saturation and increasing sensitivity^41^.

Activation-based neuronally intrinsic suppression could evolve under experimental conditions that aim to maximize recognition under conditions where objects are embedded in noise, which relates to previous studies suggesting that a general goal of adaptation is to decrease salience of recently seen stimuli or features^12,23^. This idea is also consistent with the hypothesis that adaptation enhances novelty detection by separating potentially relevant information from the prevailing input^22,36,42^. In the proposed computational model, the response strength of units for a given stimulus sequence was inversely proportional to the probability of each stimulus. Previous modeling work^43^ has suggested that adaptation effects can account for the encoding of stimulus probability in macaque IT neurons^44^. Such encoding of stimulus probability has been observed across species in somatosensory^42,45^, auditory^36,46,47^, and visual cortices^22,48,49^.

There are several limitations and possible extensions to the proposed model. We modeled fatigue as an exponential process based on the work of Bellec and colleagues^50^, therefore limiting adaptation effects to a single time scale. In reality, adaptation operates over a range of time scales from milliseconds to minutes^5^ and it has been proposed that the mechanisms can be better described by a scale-invariant power-law^51,52^. However, power-law adaptation can be approximated over a finite time interval using a sum of exponential functions^52^, and therefore the proposed model could be extended with additional exponential processes.

In the proposed model, adaptation is implemented at the level of the whole neuron. However, there exist also adaptation mechanisms that are synapse specific. Repetitive stimulation can cause short-term depression by decreasing neurotransmitter release through several molecular mechanisms (for an overview see^53^). For example, depression at thalamocortical synapses could contribute to adaptation in sensory cortex^54^.

We focused exclusively on suppressive effects of adaptation. While response attenuation is the most prevalent effect of adaptation, previous work has shown that adaptation can sometimes lead to response facilitation^55,56^. This repetition enhancement can be explained by suppressive effects that interact with normalization signals by altering the balance between excitatory and inhibitory inputs^12^. In the rat data from the oddball experiment presented here (**Fig. 2d-j**), we found a response enhancement for the rare deviant stimulus in higher visual cortex, in addition to repetition suppression for the standard stimulus^22^. It is unclear whether this enhancement arose from an interaction between adaptation effects and circuit mechanisms, or whether this observation indicates an entirely unrelated mechanism. Our proposed computational model did not capture these response enhancements, but future iterations could implement circuit level computations to examine how they interact with neuronal fatigue. Furthermore, it should be noted that, consistent with the proposed model, a deviant enhancement was not observed in macaque IT^48^.

Neuronally intrinsic mechanisms and circuit-based mechanisms of adaptation are not mutually exclusive. One common circuit-based implementation of adaptation is based on the idea of predictive coding^57^. Repetition suppression has been interpreted as a manifestation of a reduced prediction error^23,35,58,59^. This framework stresses the role of top-down modulations by internally generated perceptual expectations. However, repetition suppression can be dissociated in time from expectation effects^60^, and is not modulated by perceptual expectations induced by repetition probability in macaque IT neurons^21,48^, suggesting that suppressive effects due to expectations and due to repetitions may be caused by separate underlying mechanisms. The current results show that feed-forward effects of intrinsic neuronal mechanisms are sufficient to account for the temporal dynamics of adaptation effects and argue that a more complex top-down circuitry is not required for these effects.

Responses may also be suppressed by inhibitory connections in the circuit. Postsynaptic inhibition has been shown to contribute to adaptation in rat auditory cortex in the first 50-100 ms after a stimulus, but it did not explain adaptation at slower time scales^61^. We trained a recurrent neural network on an object recognition task that involved adaptation, which the network learned to solve using a circuit implementation. Interestingly, in contrast with the proposed fatigue model, the solution of the recurrent net was less robust to noise conditions that differed from training. While we do not exclude that a circuit implementation exists that is as robust as intrinsic fatigue in this experiment, the results do suggest that intrinsic fatigue provides a simple solution that efficiently generalizes to different conditions.

Overall, the current framework connects systems to cellular neuroscience in one comprehensive multi-scale model including an intrinsic fatigue mechanism in a deep neural network. The results demonstrate that response fatigue cascading through a feedforward hierarchical network is sufficient to explain the behavioral and neural hallmarks of visual adaptation. Intrinsic neural mechanisms may contribute substantially to the dynamics of sensory processing and perception in a temporal context.

## Methods

### Computational Models

#### Adaptation

We used the AlexNet architecture^20^ (**Fig. 1a**), with weights pre-trained on the ImageNet dataset^62^ as a model for the ventral visual stream. We implemented an exponentially decaying fatigue mechanism^50^. For each unit in every convolutional and fully connected layer (except for the decoder), we assigned an adaptation state *s_t_*, which was updated at each time step *t* based on its previous state *s*_*t*–1_ and the previous response *r*_*t*–1_ (i.e. activation after the ReLU rectification and linearization operation):

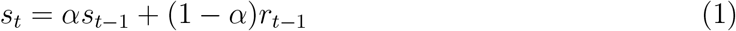

where *α* is a constant in [0,1] determining the time scale of the decay (**Fig. 1b**). This suppression state is then subtracted from the unit’s current input *x_t_* (given weights *W* and bias *b*) before applying the rectifier activation function *σ*, so that:

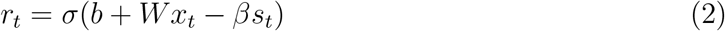

where *β* is a constant that scales the amount of suppression. These model updating rules result in an exponentially decaying response for constant input which recovers in case of no input (**Fig. 1b**), simulating a neuronal fatigue mechanism intrinsic to each individual neuron. By implementing this mechanism across discrete time steps in AlexNet, we introduced a temporal dimension to the network (**Fig. 1c**). This model was implemented using TensorFlow v1.11 in Python. Throughout the paper, we use *α* = 0.96 and *β* = 0.7 unless indicated otherwise (in **Fig. 5**, those parameters are tuned).

#### Fine-tuning

To simulate the psychophysics experiment of **Fig. 4**, we fine-tuned the pre-trained fully connected layers of AlexNet to classify high contrast (i.e. 40% as opposed to 22% in the experiment) doodles on a noisy background. We used a set of 50,000 doodle images (10,000 per category) that were different from the ones used in the experiment and fine-tuned the fully connected layers of AlexNet (without fatigue) for 5 epochs (i.e. 5 full cycles through the training images), with every epoch using a different noise background for each image. We used the Adam optimization algorithm^63^ with a learning rate of 0.001, the sparse softmax cross entropy between logits and labels cost function, a batch size of 100, and no dropout.

#### Model training

The models in **Fig. 5** were trained on the doodle classification task using TensorFlow v1.11 in Python. We used a training set of 500,000 doodle images (100,000 per category), with a separate set of 1,000 images to select hyperparameters and evaluate the loss and accuracy during training. We used the Adam optimization algorithm^63^ with a learning rate of 0.001, the sparse softmax cross entropy between logits and labels cost function, a batch size of 100, and 50% training dropout in fully connected layers. The adaptation state *β* parameters were initialized at 0 (i.e. no adaptation) and the *α* parameters were initialized using a uniform distribution ranging from 0 to 1. For the weights, we used Gaussian initialization, with the scale correction proposed by^64^. Each model was trained for 5 epochs on the training set, which was sufficient for the loss and accuracy to saturate. Generalization performance was then tested on a third independent set of 5,000 images.

### Neurophysiology

We present neurophysiological data from two previously published studies in order to compare them with the neural adaptation effects of the proposed computational model: single cell (*N* = 97) recordings from inferior temporal (IT) cortex of one macaque monkey G^21^ and multi-unit recordings from primary visual cortex (*N* = 55) and latero-intermediate visual area (*N* = 48) of three rats^22^. For methodological details about the recordings and the tasks, we refer to the original papers.

### Psychophysics

Before starting the data collection, we preregistered the study design and hypothesis on the Open Science Framework (at https://osf.io/ezj8g/).

#### Participants

A total of 17 volunteers (10 female, ages 19-50) participated in our doodle categorization experiments (**Fig. 4**). In accordance with our preregistered data exclusion rule, two male participants were excluded from analyses because we could not record eye tracking data. All subjects gave informed consent and the studies were approved by the Institutional Review Board at Children’s Hospital, Harvard Medical School.

#### Stimuli

The stimulus set consisted of hand drawn doodles of apples, cars, faces, fish, and flowers from the *Quick, Draw!* dataset^31^. We selected a total of 540 doodles (108 from each of the five categories) that were judged complete and identifiable. We lowered the contrast of each doodle image (28×28 pixels) to either 22 or 29% of the original contrast, before adding a Gaussian noise pattern (SD = 0.165 in normalized pixel values) of the same resolution. The higher contrast level (29%) was chosen as a control so that the doodle was relatively visible, was used in only one sixth of the trials, and was not included in the analyses. The average categorization performance on these high contrast trials was 74% (*SD* = 8.3%), versus 63% (*SD* = 8.9%) in the low contrast trials.

#### Experimental protocol

Participants had to fixate a cross at the center of the screen in order to start a trial. Next, an adapter image was presented (for 0.5, 2, or 4 s), followed by a blank interval (of 50, 250, or 500 ms), a test image (for 500 ms), and finally a response prompt screen. The test images were noisy doodles described in the above paragraph. The adapter image could either be: an empty frame (defined by a white square filled with the background color), the same mosaic noise pattern as the one of the subsequent test image, or a randomly generated different noise pattern (**Fig. 4**). Participants were asked to keep looking at the fixation cross, which remained visible throughout the entire trial, until they were prompted to classify the test image using keyboard keys 1-5. All images were presented at 9 × 9° from a viewing distance of approximately 52 cm on a 19 inch CRT monitor (Sony Multiscan G520, 1024 × 1280 resolution), while we continuously tracked eye movements using a video-based eye tracker (EyeLink 1000, SR Research, Canada). Trials where the root-mean-square deviation of the eye-movements exceeded 1 degree of visual angle during adapter presentation were excluded from further analyses. The experiment was controlled by custom code written in MATLAB using Psychophysics Toolbox Version 3.0^65^.

## Acknowledgements

This work was supported by Research Foundation Flanders, Belgium (fellowship of K.V.), by NIH grant R01EY026025 and by the Center for Brains, Minds and Machines, funded by NSF Science and Technology Centers Award CCF-1231216.

## Contributions

K.V. conceived the model and experiment; K.V., X.B., and G.K. designed the model and experiment; K.V. collected the data, implemented the model, and carried out analyses; K.V. and G.K. wrote the manuscript, with contributions from X.B.

## References

[1] R Addams. An account of a peculiar optical phenomenon seen after having looked at a moving body. London and Edinburgh Philosophical Magazine and Journal of Science, 5:373–374, 1834.

[2] Clarissa J. Whitmire and Garrett B. Stanley. Rapid Sensory Adaptation Redux: A Circuit Perspective. Neuron, 92(2):298–315, 2016. ISSN 08966273. doi: 10.1016/j.neuron.2016.09.046. URL http://linkinghub.elsevier.com/retrieve/pii/S0896627316306511.

[3] C Blakemore and F W Campbell. On the existence of neurones in the human visual system selectively sensitive to the orientation and size of retinal images. Journal of Physiology, 203(1):237–260, 6 1969. ISSN 0955-3002.

[4] Michael A. Webster. Visual Adaptation. Annual Review of Vision Science, 1(1): 547–567, 2015. ISSN 2374-4642. doi: 10.1146/annurev-vision-082114-035509. URL http://www.annualreviews.org/doi/10.1146/annurev-vision-082114-035509.

[5] Adam Kohn. Visual Adaptation: Physiology, Mechanisms, and Functional Benefits. Journal of Neurophysiology, 10461:3155–3164, 2007. ISSN 0022-3077. doi: 10.1152/jn.00086.2007.

[6] M. Carandini and David Ferster. A Tonic Hyperpolarization Underlying Contrast Adaptation in Cat Visual Cortex. Science, 276(5314):949–952, 1997. ISSN 00368075. doi: 10.1126/science.276.5314.949. URL http://www.sciencemag.org/cgi/doi/10.1126/science.276.5314.949.

[7] M V Sanchez-Vives, L G Nowak, and David A McCormick. Membrane mechanisms underlying contrast adaptation in cat area 17 in vivo. Journal of Neuroscience, 20(11): 4267–4285, 2000. ISSN 1529-2401. doi: 20/11/4267[pii]. URL http://www.ncbi.nlm.nih.gov/pubmed/10818163.

[8] M V Sanchez-Vives, L G Nowak, and D A McCormick. Cellular mechanisms of long-lasting adaptation in visual cortical neurons in vitro. The Journal of neuroscience: the official journal of the Society for Neuroscience, 20(11):4286–4299, 2000. ISSN 1529-2401. doi: 20/11/4286[pii].

[9] Juan M. Abolafia, R. Vergara, M. M. Arnold, R. Reig, and M. V. Sanchez-Vives. Cortical Auditory Adaptation in the Awake Rat and the Role of Potassium Currents. Cerebral Cortex, 21(5):977–990, 5 2011. ISSN 1460-2199. doi: 10.1093/cercor/bhq163. URL https://academic.oup.com/cercor/article-lookup/doi/10.1093/cercor/bhq163.

[10] Wouter De Baene and Rufin Vogels. Effects of adaptation on the stimulus selectivity of macaque inferior temporal spiking activity and local field potentials. Cerebral Cortex, 20(9):2145–65, 9 2010. ISSN 1460-2199. doi: 10.1093/cercor/bhp277. URL http://www.ncbi.nlm.nih.gov/pubmed/20038542.

[11] Neel T. Dhruv and Matteo Carandini. Cascaded Effects of Spatial Adaptation in the Early Visual System. Neuron, 81(3):529–535, 2014. ISSN 08966273. doi: 10.1016/j.neuron.2013.11.025. URL http://dx.doi.org/10.1016/j.neuron.2013.11.025.

[12] Samuel G. Solomon and Adam Kohn. Moving Sensory Adaptation beyond Suppressive Effects in Single Neurons. Current Biology, 24(20):R1012–R1022, 10 2014. ISSN 09609822. doi: 10.1016/j.cub.2014.09.001. URL http://linkinghub.elsevier.com/retrieve/pii/S0960982214011166.

[13] Charles F. Cadieu, Ha Hong, Daniel L K Yamins, Nicolas Pinto, Diego Ardila, Ethan A. Solomon, Najib J. Majaj, and James J. DiCarlo. Deep Neural Networks Rival the Representation of Primate IT Cortex for Core Visual Object Recognition. PLoS Computational Biology, 10(12), 2014. ISSN 15537358. doi: 10.1371/journal.pcbi.1003963.

[14] D. L. K. Yamins, H. Hong, C. F. Cadieu, E. A. Solomon, D. Seibert, and J. J. DiCarlo. Performance-optimized hierarchical models predict neural responses in higher visual cortex. Proceedings of the National Academy of Sciences, 111(23):8619–8624, 2014. ISSN 0027-8424. doi: 10.1073/pnas.1403112111. URL http://www.pnas.org/cgi/doi/10.1073/pnas.1403112111.

[15] Umut. Güçlü and Marcel A. J. van Gerven. Deep Neural Networks Reveal a Gradient in the Complexity of Neural Representations across the Ventral Stream. Journal of Neuroscience, 35(27):10005–10014, 7 2015. ISSN 0270-6474. doi: 10.1523/JNEUROSCI.5023-14.2015. URL http://www.jneurosci.org/cgi/doi/10.1523/JNEUROSCI.5023-14.2015http://arxiv.org/abs/1411.6422http://dx.doi.org/10.1523/JNEUROSCI.5023-14.2015.

[16] Ioannis Kalfas, Satwant Kumar, and Rufin Vogels. Shape Selectivity of Middle Superior Temporal Sulcus Body Patch Neurons. Eneuro, 4(June):0113–17, 2017. ISSN 2373-2822. doi: 10.1523/ENEURO.0113-17.2017. URL http://eneuro.sfn.org/lookup/doi/10.1523/ENEURO.0113-17.2017.

[17] Ioannis Kalfas, Kasper Vinken, and Rufin Vogels. Representations of regular and irregular shapes by deep Convolutional Neural Networks, monkey inferotemporal neurons and human judgments. PLOS Computational Biology, 14(10):e1006557, 2018. ISSN 1553-7358. doi: 10.1371/journal.pcbi.1006557. URL http://dx.plos.org/10.1371/journal.pcbi.1006557.

[18] Dean A. Pospisil, Anitha Pasupathy, and Wyeth Bair. ’Artiphysiology’ reveals V4-like shape tuning in a deep network trained for image classification. eLife, 7:1–31, 2018. ISSN 2050084X. doi: 10.7554/eLife.38242.

[19] Jonas Kubilius, Stefania Bracci, and Hans P. Op de Beeck. Deep Neural Networks as a Computational Model for Human Shape Sensitivity. PLoS Computational Biology, 12(4):1–26, 2016. ISSN 15537358. doi: 10.1371/journal.pcbi.1004896.

[20] Alex Krizhevsky, Ilya Sutskever, and Geoffrey E Hinton. ImageNet Classification with Deep Convolutional Neural Networks. Advances In Neural Information Processing Systems, pages 1–9, 2012. ISSN 10495258. doi: http://dx.doi.org/10.1016/j.protcy.2014.09.007.

[21] Kasper Vinken, Hans P. Op de Beeck, and Rufin Vogels. Face Repetition Probability Does Not Affect Repetition Suppression in Macaque Inferotemporal Cortex. The Journal of Neuroscience, 38(34):7492–7504, 8 2018. ISSN 0270-6474. doi: 10.1523/JNEUROSCI.0462-18.2018. URL http://www.jneurosci.org/lookup/doi/10.1523/JNEUROSCI.0462-18.2018.

[22] Kasper Vinken, Rufin Vogels, and Hans Op de Beeck. Recent Visual Experience Shapes Visual Processing in Rats through Stimulus-Specific Adaptation and Response Enhancement. Current Biology, 27(6):914–919, 3 2017. ISSN 09609822. doi: 10.1016/j.cub.2017.02.024. URL http://dx.doi.org/10.1016/j.cub.2017.02.024http://linkinghub.elsevier.com/retrieve/pii/S0960982217301616.

[23] Rufin Vogels. Sources of adaptation of inferior temporal cortical responses. Cortex, 80: 185–195, 7 2016. ISSN 00109452. doi: 10.1016/j.cortex.2015.08.024. URL http://dx.doi.org/10.1016/j.cortex.2015.08.024http://linkinghub.elsevier.com/retrieve/pii/S0010945215003342https://linkinghub.elsevier.com/retrieve/pii/S0010945215003342.

[24] Martin Schrimpf, Jonas Kubilius, Ha Hong, Najib J. Majaj, Rishi Rajalingham, Elias B. Issa, Kohitij Kar, Pouya Bashivan, Jonathan Prescott-Roy, Kailyn Schmidt, Daniel L. K. Yamins, and James J. DiCarlo. Brain-Score: Which Artificial Neural Network for Object Recognition is most Brain-Like? bioRxiv, page 407007, 2018. doi: 10.1101/407007. URL https://www.biorxiv.org/content/10.1101/407007v1.

[25] Hiromasa Sawamura, Guy A. Orban, and Rufin Vogels. Selectivity of neuronal adaptation does not match response selectivity: a single-cell study of the FMRI adaptation paradigm. Neuron, 49(2):307–318, 1 2006. ISSN 0896-6273. doi: 10.1016/j.neuron.2005.11.028. URL http://www.ncbi.nlm.nih.gov/pubmed/16423703.

[26] J. J. Gibson and Minnie Radner. Adaptation, after-effect and contrast in the perception of tilted lines. I. Quantitative studies. Journal of Experimental Psychology, 20(5):453–467, 1937. ISSN 0022-1015. doi: 10.1037/h0059826. URL http://doi.apa.org/getdoi.cfm?doi=10.1037/h0059826.

[27] JP Frisby and JV Stone. Seeing Aftereffects: The Psychologist’s Microelectrode. In Seeing: The computational approach to biological vision, pages 75–110. MIT Press, Cambridge, MA, 2 edition, 2010. ISBN 0262514273.

[28] Felix A Wichmann and N Jeremy Hill. The psychometric function: I. Fitting, sampling, and goodness of fit. Perception & Psychophysics, 63(8):1293–1313, 11 2001. ISSN 0031-5117. doi: 10.3758/BF03194544. URL http://www.springerlink.com/index/10.3758/BF03194544.

[29] Michael A. Webster, Daniel Kaping, Yoko Mizokami, and Paul Duhamel. Adaptation to natural facial categories. Nature, 428(6982):557–561, 4 2004. ISSN 0028-0836. doi: 10.1038/nature02420. URL http://www.nature.com/doifinder/10.1038/nature02420.

[30] Lisa M DeBruine. Webmorph (Version v0.0.0.9001), 2019.

[31] Google Creative Lab. The Quick, Draw! Dataset. GitHub repository https://github.com/googlecreativelab/quickdraw-dataset, 2019.

[32] Colin W G Clifford, Michael A. Webster, Garrett B. Stanley, Alan A. Stocker, Adam Kohn, Tatyana O. Sharpee, and Odelia Schwartz. Visual adaptation: Neural, psychological and computational aspects. Vision Research, 47(25):3125–3131, 2007. ISSN 00426989. doi: 10.1016/j.visres.2007.08.023.

[33] Hanlin Tang, Martin Schrimpf, William Lotter, Charlotte Moerman, Ana Paredes, Josue Ortega Caro, Walter Hardesty, David Cox, and Gabriel Kreiman. Recurrent computations for visual pattern completion. Proceedings of the National Academy of Sciences, page 201719397, 2018. ISSN 0027-8424. doi: 10.1073/pnas.1719397115. URL http://www.pnas.org/lookup/doi/10.1073/pnas.1719397115.

[34] Kohitij Kar, Jonas Kubilius, Kailyn Schmidt, Elias B Issa, and James J. DiCarlo. Evidence that recurrent circuits are critical to the ventral stream’s execution of core object recognition behavior. Nature Neuroscience, page 354753, 4 2019. ISSN 1097-6256. doi: 10.1038/s41593-019-0392-5. URL http://dx.doi.org/10.1038/s41593-019-0392-5http://www.nature.com/articles/s41593-019-0392-5https://www.biorxiv.org/content/10.1101/354753v1.

[35] William Lotter, Gabriel Kreiman, and David Cox. A neural network trained to predict future video frames mimics critical properties of biological neuronal responses and perception. arXiv preprint, pages 1–18, 5 2018. ISSN 01677322. doi: 10.1016/j.molliq.2011.09.016. URL https://linkinghub.elsevier.com/retrieve/pii/S0167732211003096http://arxiv.org/abs/1805.10734.

[36] Nachum Ulanovsky, Liora Las, and Israel Nelken. Processing of low-probability sounds by cortical neurons. Nature neuroscience, 6(4):391–8, 4 2003. ISSN 1097-6256. doi: 10.1038/nn1032. URL http://www.ncbi.nlm.nih.gov/pubmed/12652303.

[37] Dzmitry A. Kaliukhovich and Hans Op de Beeck. Hierarchical stimulus processing in rodent primary and lateral visual cortex as assessed through neuronal selectivity and repetition suppression. Journal of Neurophysiology, 120(3):926–941, 2018. ISSN 0022-3077. doi: 10.1152/jn.00673.2017.

[38] Javier Nieto-Diego and Manuel S. Malmierca. Topographic Distribution of Stimulus-Specific Adaptation across Auditory Cortical Fields in the Anesthetized Rat. PLOS Biology, 14(3):e1002397, 3 2016. ISSN 1545-7885. doi: 10.1371/journal.pbio.1002397. URL http://dx.plos.org/10.1371/journal.pbio.1002397.

[39] S Laughlin. Energy as a constraint on the coding and processing of sensory information. Current Opinion in Neurobiology, 11(4):475–480, 8 2001. ISSN 09594388. doi: 10.1016/S0959-4388(00)00237-3. URL https://linkinghub.elsevier.com/retrieve/pii/S0959438800002373.

[40] Peter Lennie. The Cost of Cortical Computation. Current Biology, 13(6):493–497, 3 2003. ISSN 09609822. doi: 10.1016/S0960-9822(03)00135-0. URL https://linkinghub.elsevier.com/retrieve/pii/S0960982203001350.

[41] Michael A. Webster, John S. Werner, and David J. Field. Adaptation and the Phenomenology of Perception. In Fitting the Mind to the WorldAdaptation and After-Effects in High-Level Vision, chapter 10, pages 241–278. Oxford University Press, 5 2005. ISBN 9780199251841. doi: 10.1093/acprof:oso/9780198529699.003.0010. URL http://www.oxfordscholarship.com/view/10.1093/acprof:oso/9780198529699.001.0001/acprof-9780198529699-chapter-10.

[42] Simon Musall, Florent Haiss, Bruno Weber, and Wolfger von der Behrens. Deviant Processing in the Primary Somatosensory Cortex. Cerebral Cortex, page bhv283, 2015. ISSN 1047-3211. doi: 10.1093/cercor/bhv283. URL http://www.cercor.oxfordjournals.org/lookup/doi/10.1093/cercor/bhv283.

[43] Kasper Vinken and Rufin Vogels. Adaptation can explain evidence for encoding of probabilistic information in macaque inferior temporal cortex. Current Biology, 27(22):R1210–R1212, 11 2017. ISSN 09609822. doi: 10.1016/j.cub.2017.09.018. URL http://linkinghub.elsevier.com/retrieve/pii/S096098221731182Xhttps://linkinghub.elsevier.com/retrieve/pii/S096098221731182X.

[44] Andrew H. Bell, Christopher Summerfield, Elyse L. Morin, Nicholas J. Malecek, and Leslie G. Ungerleider. Encoding of Stimulus Probability in Macaque Inferior Temporal Cortex. Current Biology, 26(17):2280–2290, 9 2016. ISSN 09609822. doi: 10.1016/j.cub.2016.07.007. URL http://dx.doi.org/10.1016/j.cub.2016.07.007http://linkinghub.elsevier.com/retrieve/pii/S0960982216307588.

[45] Simon Musall, Wolfger von der Behrens, Johannes M Mayrhofer, Bruno Weber, Fritjof Helmchen, and Florent Haiss. Tactile frequency discrimination is enhanced by circumventing neocortical adaptation. Nature neuroscience, 17(11):1567–73, 2014. ISSN 1546-1726. doi: 10.1038/nn.3821. URL http://www.ncbi.nlm.nih.gov/pubmed/25242306.

[46] Y. I. Fishman and M. Steinschneider. Searching for the Mismatch Negativity in Primary Auditory Cortex of the Awake Monkey: Deviance Detection or Stimulus Specific Adaptation? Journal of Neuroscience, 32(45):15747–15758, 11 2012. ISSN 0270-6474. doi: 10.1523/JNEUROSCI.2835-12.2012. URL http://www.jneurosci.org/cgi/doi/10.1523/JNEUROSCI.2835-12.2012.

[47] Brandon J Farley, Michael C Quirk, James J Doherty, and Edward P Christian. Stimulus-specific adaptation in auditory cortex is an NMDA-independent process distinct from the sensory novelty encoded by the mismatch negativity. The Journal of neuroscience: the official journal of the Society for Neuroscience, 30(49):16475–84, 12 2010. ISSN 1529-2401. doi: 10.1523/JNEUROSCI.2793-10.2010. URL http://www.ncbi.nlm.nih.gov/pubmed/21147987.

[48] D. A. Kaliukhovich and R. Vogels. Neurons in Macaque Inferior Temporal Cortex Show No Surprise Response to Deviants in Visual Oddball Sequences. Journal of Neuroscience, 34(38):12801–12815, 9 2014. ISSN 0270-6474. doi: 10.1523/JNEUROSCI.2154-14.2014. URL http://www.jneurosci.org/cgi/doi/10.1523/JNEUROSCI.2154-14.2014.

[49] Jordan P. Hamm and Rafael Yuste. Somatostatin Interneurons Control a Key Component of Mismatch Negativity in Mouse Visual Cortex. Cell Reports, 16(3): 597–604, 2016. ISSN 22111247. doi: 10.1016/j.celrep.2016.06.037. URL http://dx.doi.org/10.1016/j.celrep.2016.06.037.

[50] Guillaume Bellec, Darjan Salaj, Anand Subramoney, Robert Legenstein, and Wolfgang Maass. Long short-term memory and learning-to-learn in networks of spiking neurons. Conference on Neural Information Processing Systems, 3 2018. doi: arXiv:1803.09574. URL http://arxiv.org/abs/1803.09574.

[51] Barry Wark, Brian Nils Lundstrom, and Adrienne Fairhall. Sensory adaptation. Current Opinion in Neurobiology, 17(4):423–429, 2007. ISSN 09594388. doi: 10.1016/j.conb.2007.07.001.

[52] Patrick J Drew and L F Abbott. Models and Properties of Power-Law Adaptation in Neural Systems. Journal of Neurophysiology, 96(2):826–833, 8 2006. ISSN 0022-3077. doi: 10.1152/jn.00134.2006. URL http://www.physiology.org/doi/10.1152/jn.00134.2006.

[53] Diasynou Fioravante and Wade G. Regehr. Short-term forms of presynaptic plasticity. Current Opinion in Neurobiology, 21(2):269–274, 2011. ISSN 09594388. doi: 10.1016/j.conb.2011.02.003. URL http://dx.doi.org/10.1016/j.conb.2011.02.003.

[54] Sooyoung Chung, Xiangrui Li, and Sacha B. Nelson. Short-term depression at thalamocortical synapses contributes to rapid adaptation of cortical sensory responses in vivo. Neuron, 34(3):437–446, 2002. ISSN 08966273. doi: 10.1016/S0896-6273(02)00659-1.

[55] Stephanie C Wissig and Adam Kohn. The influence of surround suppression on adaptation effects in primary visual cortex. Journal of Neurophysiology, 107(12): 3370–3384, 6 2012. ISSN 1522-1598. doi: 10.1152/jn.00739.2011. URL http://www.ncbi.nlm.nih.gov/pubmed/22423001.

[56] Dzmitry A. Kaliukhovich and Rufin Vogels. Divisive Normalization Predicts Adaptation-Induced Response Changes in Macaque Inferior Temporal Cortex. The Journal of Neuroscience, 36(22):6116–6128, 2016. ISSN 0270-6474. doi: 10.1523/JNEUROSCI.2011-15.2016. URL http://www.jneurosci.org/lookup/doi/10.1523/JNEUROSCI.2011-15.2016.

[57] Rajesh P N Rao and Dana H Ballard. Predictive coding in the visual cortex: a functional interpretation of some extra-classical receptive-field effects. Nature neuroscience, 2(1):79–87, 1999. ISSN 1097-6256. doi: 10.1038/4580. URL 10.1038/4580%5Cnhttp://www.nature.com/neuro/journal/v2/n1/abs/nn0199_79.html.

[58] Karl Friston. A theory of cortical responses. Philosophical transactions of the Royal Society of London B, 360(1456):815–36, 4 2005. ISSN 0962-8436. doi: 10.1098/rstb.2005.1622. URL http://www.pubmedcentral.nih.gov/articlerender.fcgi?artid=1569488&tool=pmcentrez&rendertype=abstracthttp://www.ncbi.nlm.nih.gov/pubmed/15937014http://www.pubmedcentral.nih.gov/articlerender.fcgi?artid=PMC1569488.

[59] Christopher Summerfield, Emily H Trittschuh, Jim M Monti, M Marsel Mesulam, and Tobias Egner. Neural repetition suppression reflects fulfilled perceptual expectations. Nature Neuroscience, 11(9):1004–1006, 9 2008. ISSN 1546-1726. doi: 10.1038/nn.2163. URL http://www.pubmedcentral.nih.gov/articlerender.fcgi?artid=2747248&tool=pmcentrez&rendertype=abstract.

[60] Ana Todorovic and Floris P de Lange. Repetition suppression and expectation suppression are dissociable in time in early auditory evoked fields. The Journal of Neuroscience, 32(39):13389–13395, 9 2012. ISSN 1529-2401. doi: 10.1523/JNEUROSCI.2227-12.2012. URL http://www.ncbi.nlm.nih.gov/pubmed/23015429.

[61] Michael Wehr and Anthony M. Zador. Synaptic mechanisms of forward suppression in rat auditory cortex. Neuron, 47(3):437–445, 2005. ISSN 08966273. doi: 10.1016/j.neuron.2005.06.009.

[62] Olga Russakovsky, Jia Deng, Hao Su, Jonathan Krause, Sanjeev Satheesh, Sean Ma, Zhiheng Huang, Andrej Karpathy, Aditya Khosla, Michael Bernstein, Alexander C. Berg, and Li Fei-Fei. ImageNet Large Scale Visual Recognition Challenge. International Journal of Computer Vision, 115(3):211–252, 2015. ISSN 15731405. doi: 10.1007/s11263-015-0816-y.

[63] Diederik P. Kingma and Jimmy Ba. Adam: A Method for Stochastic Optimization. Proceedings of the 3rd International Conference on Learning Representations (ICLR), pages 1–15, 12 2014. URL http://arxiv.org/abs/1412.6980.

[64] Xavier Glorot and Yoshua Bengio. Understanding the difficulty of training deep feedforward neural networks. Proceedings of the 13th International Conference On Artificial Intelligence and Statistics, 9:249–256, 2010.

[65] David H. Brainard. The Psychophysics Toolbox. Spatial Vision, 10(4):433–436, 1997. ISSN 01691015. doi: 10.1163/156856897X00357.

